# Identifying genomic markers associated with female re-mating rate in *Drosophila pseudoobscura* by replicated bulk segregant analysis

**DOI:** 10.1101/2020.04.20.049940

**Authors:** R Axel W Wiberg, Tom A R Price, Nina Wedell, Michael G Ritchie

**Author notes:** Department of Environmental Sciences, Zoological Institute, University of Basel, Vesalgasse 1, 4051 Basel, Switzerland (R Axel W Wiberg).

## Abstract

Identifying loci associated with a phenotype is a critical step in many evolutionary studies. Most methods require large sample sizes or breeding designs that can be prohibitively difficult. Here we apply a rarely used approach to identify SNP loci associated with a complex phenotype. We mate siblings from isofemale lines isolate genotypes from three wild populations. After phenotyping we perform whole genome sequencing of isofemale lines from the extremes of the phenotypic distribution of each population and identify SNPs that are consistently fixed for alternative alleles across line pairs. The focal phenotype is female remating rate in the fly *Drosophila pseudoobscura,* defined as the willingness of a female to mate with a second male after her first mating. This is an integral part of mating system evolution, sexual selection and sexual conflict, and is a quantitative polygenic trait.

About 200 SNPs are consistently fixed for alternate alleles in the three pairs of isofemale lines. We use different simulation approaches to explore how many SNPs would be expected to be fixed. We find the surprising result that we uncover *fewer* observed fixed SNPs than are expected by either simulation approach. We also complete functional analyses of these SNPs. Many lie near genes or regulatory regions known to be involved in *Drosophila* courtship and mating behaviours, and some have previously been associated with re-mating rates in Genome-Wide Association Studies. Given the small sample size, these results should be treated with caution. Nevertheless, this study suggests that even from a relatively small number of isofemale lines established from wild populations, it is possible to identify candidate loci potentially associated with a complex quantitative trait. However, further work is required to understand modelling the expected distribution of differences.

## INTRODUCTION

Identifying the genetic basis of quantitative traits continues to be an important goal in evolutionary biology, and multiple approaches are available (Boake et al., 2002; Stapley et al., 2010; Hoban et al., 2016). Quantitative Trait Locus (QTL) mapping relies on associating genomic markers with a trait and many studies have successfully identified QTL for traits important in adaptation (e.g. Colosimo et al., 2004; Kronforst et al., 2006). However, such studies are laborious, require inbred lines or extensive pedigrees, typically have low resolution, and loci with small effects can be missed entirely (Rockman, 2011; Travisano & Shaw, 2012). As genome-wide sequencing and genotyping at large numbers of loci has become increasingly accessible, Genome-Wide Association Studies (GWAS) have become a more popular approach (Stapley et al., 2010; Hoban et al., 2016) and may have a greater genomic resolution because they typically involve many more markers, but consequently can often require very large sample sizes at considerable expense.

Innovations for both model and non-model study systems have been accumulating (Schlötterer et al., 2014; Schneeberger, 2014). For example, pooled-sequencing (pool-seq) can identify accurate gene frequency data for large numbers of individuals for a fraction of the price of multiple individual whole genome sequences (Schlötterer et al., 2014). While haplotype information is lost, this approach has proven successful in detecting allele frequency differences between natural populations (e.g. Lamichhaney et al., 2012; Bergland et al., 2014; Bhattacharjee et al., 2016; Chen et al., 2016) and between treatment groups in experimental evolution studies (e.g. Burke et al., 2010; Orozco-terWengel et al., 2012; Kofler & Schlötterer, 2014; Schlötterer et al., 2015). Bulk segregant analysis (BSA) is an approach which leverages the extremes of the distribution of a quantitative trait in order to identify loci associated with this variation (Magwene et al., 2011). Individuals from the upper and lower tails of the distribution are sequenced. Genomic regions which have no effect on the trait should show random allele frequency differences between the two extremes. Meanwhile, regions containing loci influencing the trait of interest should show more consistent differences in allele frequencies (Magwene et al., 2011), and these should be more consistent across multiple comparisons of extreme phenotype pools. Using extreme phenotypes has also been extended to GWAS, Extreme-phenotype GWAS (XP-GWAS; Yang et al., 2015). Causative alleles are likely to be enriched in pools of extreme phenotypes, and greater resolution is achieved by using more markers (e.g. by using SNP-chips, or whole-genome sequencing; Yang et al., 2015).

However, identifying individuals with the genotypes for extreme phenotypes can be difficult, as environmental effects and chance may also contribute to extremes. One solution is to use inbred lines. In this technique a single female is collected in the wild, and her offspring are inbred via sib-sib matings for several generations to produce inbred lines where all individuals are almost genetically identical (David et al., 2005). Each inbred line effectively captures a wild genotype, creating a suite of lines of genetically identical individuals in the laboratory. Isofemale lines can then be reliably phenotyped across many individuals and extreme lines are candidates for genotyping. Hence isofemale or inbred lines are a staple of research and collections of lines exist as a public resource for several species including *Arabidopsis thaliana* (The 1001 Genomes Consortium 2016) and *Drosophila melanogaster* (Mackay et al., 2012; Huang et al., 2014). GWAS or QTL studies can be carried out directly on these inbred lines (e.g. Montgomery et al., 2014; Ivanov et al., 2015; Gaertner et al., 2015) but again resolution can be severely limited by sample size.

Although costs of sequencing have fallen dramatically it can still be relatively high for many researchers and the feasibility of a method often depends on the species. Methods that can reliably identify associated loci while reducing the sample sizes and sequencing effort are particularly useful for non-model systems. In this study we test a method of identifying SNP markers with a complex polygenic behavioural trait in isofemale lines recently collected from the wild. We use three pairs of lines from the extremes of the phenotypic distribution of three different populations (figure 1). Pool-seq is used to identify fixed differences between these three pairs of lines, which is analogous to three replicated bulk segregant analyses (or XP-GWAS; see above) replicated across populations rather than within the extreme phenotype pools. Any single pairwise comparison will produce fixed differences between lines, many of which will be by chance. However, combining multiple pairwise comparisons from different populations (with different demographic and evolutionary histories) to identify only sites with consistently fixed differences should reduce spurious chance differences (figure 1). This should be especially true for species with large natural effective population sizes and genomes shaped by many rounds of recombination in the wild. The number of fixed differences identified is compared to expectations from population genetic simulations. Finally, the chromosomal positions of fixed differences and the linked genes or regulatory regions can also be identified and any known association with the phenotype checked.

**Figure 1.**
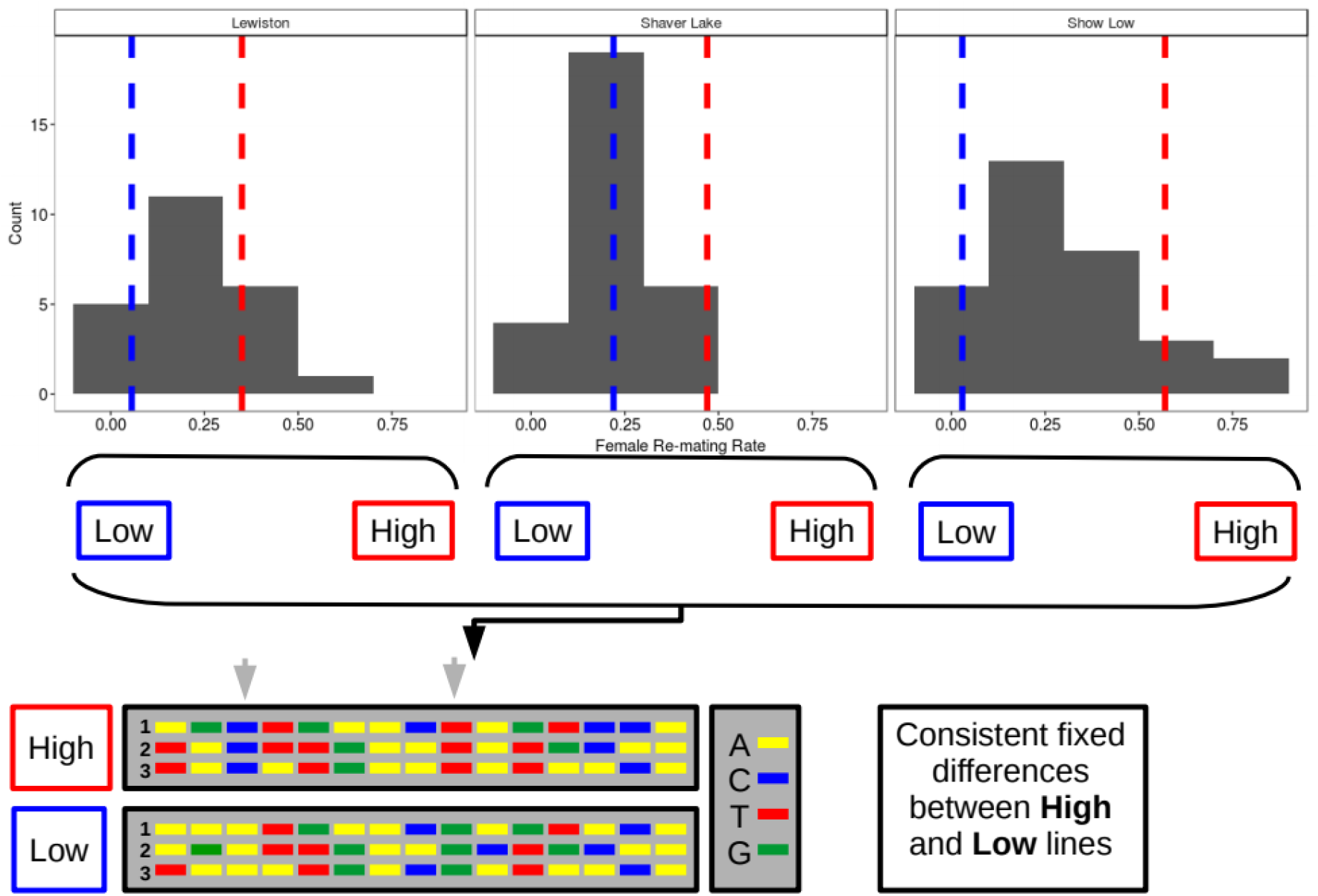
Cartoon representation of the method. Isofemale lines from different populations show variation in female re-mating rates. Histograms show the variation in re-mating rate among lines from Lewston, Shaver Lake, and Show Low (see also table S1). Lines from the tails of these distributions represent “High” and “Low” re-mating rates (vertical blue and red dashed lines). Pool-seq data allows the identification of SNPs that show fixed differences (grey arrows) between High and Low lines across all pairwise comparisons.

The trait we examine is female re-mating rate in *D. pseudoobscura*. Female mating rate is an important component of sexual selection and conflict (Pizzari & Wedell, 2013; Snook 2014) and varies widely between species and natural populations in this species (Price et al., 2014). This variation probably has a strong genetic component. Re-mating rates of wild females, estimated from the number of sires represented among the offspring, are correlated with the latency to re-mate among daughters in the laboratory (Price *et al.,* 2011). Granddaughters of females caught in populations where re-mating rates are high tend to have higher re-mating rates, and shorter latencies to re-mate, than grand-daughters of females caught in populations with low re-mating rates (Price *et al.,* 2014). In addition, the female re-mating rate can evolve rapidly in experimental evolution studies (Price et al., 2008), and males have little ability to suppress re-mating by females (Price et al 2010).

Understanding the causes and consequences of female re-mating rates in *D. pseudoobscura* will require an improved picture of the genetic basis of the trait. Most work on the genetics of female re-mating has previously been done with *D. melanogaster* (e.g. McGraw et al., 2004, 2008; Swanson et al., 2004; Mackay et al., 2005; Giardina et al., 2011; Giardina, 2015). These have generally been mutant screens or QTL studies and have identified potential candidate genes for female re-mating rate. For example, olfactory receptor genes and odorant binding proteins are known to be upregulated in females as a response to mating (McGraw et al., 2004, 2008) and are associated with re-mating among female lines that vary in re-mating rate (Giardina et al., 2011). Also implicated are many genes involved in the seminal fluid cocktail passed by the males (Ram & Wolfner, 2007). Perhaps the most well studied is “sex peptide,” a male accessory gland protein which interacts with female receptors in the reproductive tract and induces a post-mating response in females of many *Drosophila* species (Chapman *et al.,* 2003; Yapici *et al.,* 2008; Tsuda *et al.,* 2015), part of which is a reduced willingness to re-mate (Ram & Wolfner, 2007). Other accessory gland proteins may also have similar functions and are therefore prime candidates for genes affecting female remating rates. These genes are known to evolve rapidly in *Drosophila* (Haerty *et al.,* 2007), function differently in different lineages (Tsuda *et al.,* 2015) and some have undergone duplications or losses in different lineages (Tsuda & Aigaki, 2016).

We identify SNPs that are fixed between the extreme lines and use a variety of different parameterisations of population genomic simulations to assess the number expected. Surprisingly, we find that we identify *fewer* observed fixed SNPs than are expected by our simulation approaches. Some of these are associated with genes known to be involved in *Drosophila* courtship and mating behaviours and have previously been associated with female re-mating rate. We suggest that even from a relatively small number of isofemale lines it is possible to identify candidate loci associated with a complex quantitative trait, but identifying the numbers expected is complex.

## METHODS

### Sample Collection

Samples were collected from three populations; Show Low, Arizona (34° 07’ 3’’N, 110° 07’ 37’’W); Lewistown, Montana (47° 04’ 47’’N, 109° 16’ 53’’W); and Shaver Lake, California (37° 8’ 50.64’’ N, 119° 18’ 6.336’’ W) (Price et al., 2014). Isofemale lines were set up from each population. Briefly, offspring from each wild caught female were inbred single sib-sib paired matings for 8 generations. Flies were then maintained in family groups for ~40 further generations in the lab. Female re-mating rate was determined in 2013, and subsequently reconfirmed in 2017. Re-mating rates were quantified as follows; Virgin females were collected and stored in single sex groups of 10 individuals. At three days old, females were moved to individual vials and at 4 days old, each was presented with a four-day old stock male and mated. Four days later at 8 days old each female was presented with a second four-day old stock male, and observed for two hours. The re-mating rate was estimated as the proportion of females re-mating in this trial. For the current study, 6 isofemale lines, two per population, were chosen from the extremes of the distribution of female re-mating rates. Summary statistics for the populations and isofemale lines are shown in table S1. Isofemale lines are not perfect matches in the rates of female re-mating (table S1, figure 1) due to some lines being unavailable by the time of sequencing; rather, they were the most extreme available.

### Sequencing and Mapping

Sequencing was carried out at the NBAF sequencing facility at the Center for Genomic Research (CGR) at the University of Liverpool. Samples were sequenced using a “pool-seq” approach (Schlötterer et al., 2014). For each isoline, 40 females were pooled and DNA extracted using a standard phenol-chloroform extraction protocol. Four libraries were run on a single Illumina HiSeq lane and sequenced to ~40x coverage. Empirical coverage statistics and the number of reads generated as well as quality metrics are shown in table S2.

Further quality control by trimming and filtering low quality reads was performed using Trimmomatic v. 0.32 (Bolger et al., 2014). Reads were clipped if the base quality fell below Q = 20 and reads shorter than 20 bp were discarded. BWA mem (Li et al., 2009; Li, 2013) was used to map reads to the *D. pseudoobscura* reference genome (release 3.1, February 2013) obtained from FlyBase (dos Santos et al., 2014). Duplicate reads were removed using samtools v. 1.2 (Li et al., 2009) and re-alignment around indels was carried out with Picard (v. 2.14.1; Broad Institute) GATK v. 3.3 (McKenna et al., 2010; DePristo et al., 2011). Bedtools v. 2.22.1 (Quinlan & Hall, 2010) was used to calculate various genome-wide statistics (e.g. coverage) throughout the genome. Summary statistics of the mapping step are shown in table 1. SNPs and allele frequencies were called with samtools v. 1.2 (Li et al., 2009) and PoPoolation2 (v. 1.201; Kofler et al., 2011). See the supplementary materials for a full pipeline of trimming, mapping and SNP calling steps along with the command line parameters used.

Due to the incompleteness of the *D. pseudoobscura* genome assembly, chromosome 4 and the X chromosome arms are split into 4 and 8 groups respectively. Also, several unmapped and fragmented scaffolds are present in the genome (~17% of the genome). Unless otherwise stated, the subsequent analyses are performed on the chromosomal groups and the unknown or unplaced scaffolds ignored. Coverage across samples is fairly consistent (figure S1 and table S1), SLOB7, SHAA10 and SHAC1 samples have higher average coverage. To avoid any confounding effects of large differences in coverage the .bam files for SLOB7, SHAA10 and SHAC1 are sub-sampled to contain 47 million alignments, corresponding to the mean across all other samples. Only biallelic SNPs with a coverage greater than 17 and lower than 59 (corresponding to the 10^th^ and 90^th^ percentiles of the aggregate coverage distribution respectively; figure S1) are considered for further analysis in order to avoid spurious SNPs from high coverage (duplicated/repetitive regions) and low coverage (poorly sequenced) genomic regions. If any sample did not meet these requirements the SNP is excluded. This left 3,709,701 SNPs for further analysis.

### Identifying Candidate SNPs

To identify SNP variants associated with female re-mating rate we first identified all SNPs that were consistently fixed for the same alternative alleles in high and low re-mating rate lines (hereafter “fixed SNPs”). Pairwise comparisons were performed between lines that come from the same population, thus there were three pairwise comparisons. Because the isofemale lines are inbred and the sample size was relatively small (three high and three low re-mating rate lines), it is possible that many of these fixed differences might be due to chance. Two different simulations were run to obtain an empirical null distribution of the number of consistently fixed differences expected by chance.

The first simulation approach considers an ancestral population at mutation-drift equilibrium in which the distribution of allele frequencies is described by a beta-binomial distribution B(α, β) (Charlesworth & Charlesworth, 2008), where the α and β shape parameters which describe the distribution are given by:

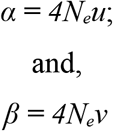

where *Ne* is the effective population size and *u* and *v* are the mutation and back-mutation rates respectively (Charlesworth & Charlesworth, 2008).

Isofemale lines can be simulated from this ancestral population by drawing allele frequencies from this distribution and assuming that these are at Hardy-Weinberg equilibrium;

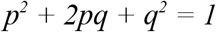

Isofemale lines are then taken to have an allele frequency of 1, 0, or 0.5 for the two homozygous genotypes and the heterozygous genotypes respectively. The inbreeding design described above (8 generations of full-sib inbreeding followed by 40 generations of maintenance in the lab) was then simulated by 8 rounds of binomial sampling and a following 40 rounds of binomial sampling with sample sizes of 4 and 80 respectively (this assumes an *N_e_* of 40 for isofemale lines throughout the 40 generations of lab maintenance). Finally, the proportion of times the allele frequency difference between pairs of isofemale lines is 1 across all *n* pairs of isofemale lines can be computed to derive a theoretical neutral distribution of such fixations.

To fully parameterise this simulation, values of *N_e_* and *u* were obtained from the literature. Several estimates of *N_e_* for *D. pseudoobscura* have been reported. Noor et al., (2000) estimate between 141,000 and 512,000 from microsatellite data while Jensen & Bachtrog (2011) give an estimate of 4.5×10^6^ from genome-wide SNP data. Meanwhile, estimates for species with similar distributions range from 2×10^6^ *(Heliconius melpomene;* Keightley et al., 2015) to 1.4×10^6^ (*D. melanogaster;* Keightley et al., 2014). The simulations were thus run over the range of *N_e_* from 1×10^6^ to 4×10^6^. Estimates of mutation rates also vary. For species similar to *D. pseudoobscura* the mutation rate is in the range 1×10^-9^ to 8.4×10^-9^ in *D. melanogaster* (Haag-Liautard et al., 2007; Keightley et al., 2014) and 2.9×10^-9^ in *H. melpomene* (Keightley et al., 2015). Because only the compound value *4N_e_μ* is relevant to these simulations we used the possible values of *4N_e_μ* from all combinations of the values from the literature above.

A second approach to assessing the expected number of consistently fixed differences is similar to a bootstrapping approach. Instead of a hypothetical ancestral allele frequency distribution this uses the empirical distribution of allele frequencies in the pool-seq data from each isofemale line. Because the distributions were very similar across all samples (figure S2), the aggregate distribution (summing counts in each frequency bin across isofemale line samples) was used (figure S2). Allele frequencies (*p*) were drawn from this empirical distribution. In this case genotypes were not assumed to be at HWE and were instead made by simply drawing two alleles at random from a binomial distribution where the probabilities were given by *p*. Thus, genotypes are either 1/1, 0/0 or 1/0 for each isofemale line with the probability of drawing a “1” being set by *p*. Isofemale lines are then given allele frequencies of 1, 0, or 0.5. Finally, the proportion of times consistent differences of 1 (i.e. fixed differences) across all pairs of isofemale lines are seen is computed. The proportion of consistent fixations was calculated from 1,000, 10,000, 100,000, and 1,000,000 SNPs drawn from the empirical distribution across *n* = 3 pairs of isofemale lines. A distribution of the proportion of consistent fixations was simulated from 100 runs for each number of SNPs. This bootstrapping approach should be conservative because there is very little variation in allele frequencies in the empirical distribution with most sites being fixed for either allele (figure S2), thus there is a high probability of individuals being homozygotes but a good chance of being homozygous for the minor allele as well as the major allele (figure S2).

### Functional Analysis

#### Gene Ontology and Phenotype Enrichment

For the identified SNPs a functional analysis was carried out by Gene Ontology (GO) term and phenotypic class enrichment analysis. These analyses rely on GO term and phenotypic associations with annotated genes in *D. melanogaster.* Thus, *D. melanogaster* GO terms were downloaded via FuncAssociate (v2.0; Berriz et al., 2009). The *D. pseudoobscura* annotated genes were converted to *D. melanogaster* orthologs, where duplicates were found they were re-labelled in the annotation and the duplicate ID was added to the GO term dataset.

GO term enrichment analysis was performed for each SNP in GOwinda (v1.12; Kofler & Schlötterer, 2012). The SNP was considered “genic” if it occurred within 1Mb up- or downstream of an annotated gene. GOwinda was run with default parameters and the empirical null distribution of gene abundance distribution obtained by 10,000 simulations.

ModPhEA (Weng & Liao, 2017) was used to carry out phenotype enrichment analysis. First SNPs were associated with a gene by identifying the closest gene within 1Mb to each fixed SNP. The set of all unique genes was submitted to modPhEA to test for an association with any phenotypic classes from the *Drosophila melanogaster* phenotypic terms. Phenotypic classes from levels 2 through 7 of the hierarchy were used to strike a balance between phenotypic detail and statistical power. A distance of up to 1Mb in GO term and phenotype enrichment analysis is justified on the basis that regulatory regions are frequently mapped to distances of ~5kb (Werner et al., 2010), ~20kb (Chan et al., 2010), and up to 1 Mb up- or downstream from a target gene (e.g. Maston et al., 2006; Pennacchio et al., 2013).

#### TF Motifs

Finally, a transcription factor (TF) motif enrichment analysis was performed with the AME routine (McLeay & Bailey 2010) from the MEME package (v. 4.10.2; Bailey et al., 2009). This tool takes a set of short DNA sequences and compares them to a database of known TF binding motifs (Fly Factor Survey; http://mccb.umassmed.edu/ffs/) to determine if any are overrepresented among the sequences. The sequence extending 30bp up- and downstream of each fixed SNP was extracted from the genome. This region is large enough to accommodate even the larger TF binding motifs but small enough that the focal SNP could conceivably be within the active region of the motif. The sequences around all SNPs were used as a control set in the AME analysis. The motif database consisted of 656 TF motifs. See the supplementary materials for a description of the pipeline.

## RESULTS

### Identifying Candidate SNPs

Mapping results are given in table S1 and figure S1. Sub-sampling of the SLOB7, SHAC1 and SHAA10 samples was carried out to equalise the coverage across samples. The majority of reads were mapped unambiguously and paired with forward and reverse reads mapping to the same scaffold (table S1). In total, 2,872,064 SNPs passed quality control. Out of these, 193 SNPs (0.007%) were consistently fixed for the same alleles in the high re-mating lines compared to the low re-mating rate lines across all pairwise comparisons (hereafter “fixed SNPs”).

Simulations of a population at mutation-drift equilibrium suggest that 193 SNPs (0.007%) is many fewer than would be expected by chance (~1% for *n* = 3 isofemale line pairs; figure 2 A and B). The proportion of consistently fixed SNPs increases with *N_e_* and decreases with increasing numbers of sampled isofemale lines (figure 2 A and B). Differences in *4N_e_μ* do not seem to have much of an effect on the number of fixed SNPs expected except at low *n*. For *n* = 3, the 95^th^ percentile of the distribution of expected proportions of fixed differences is 1.02% (figure 2 A and B), suggesting that 0.007% is a much lower proportion of fixed differences than expected by chance. Similarly, bootstrap sampling of SNPs and allele frequencies from the empirical distribution suggests that the proportion of consistently fixed differences expected by chance is considerably higher, around 0.4%, regardless of the number of SNPs sampled (1,000, 10,000, 100,000, or 1,000,000; figure 2C). The range of the 95^th^ percentile of the distribution of expected fixed differences is between 0.9 and 0.41% (data not shown).

**Figure 2.**
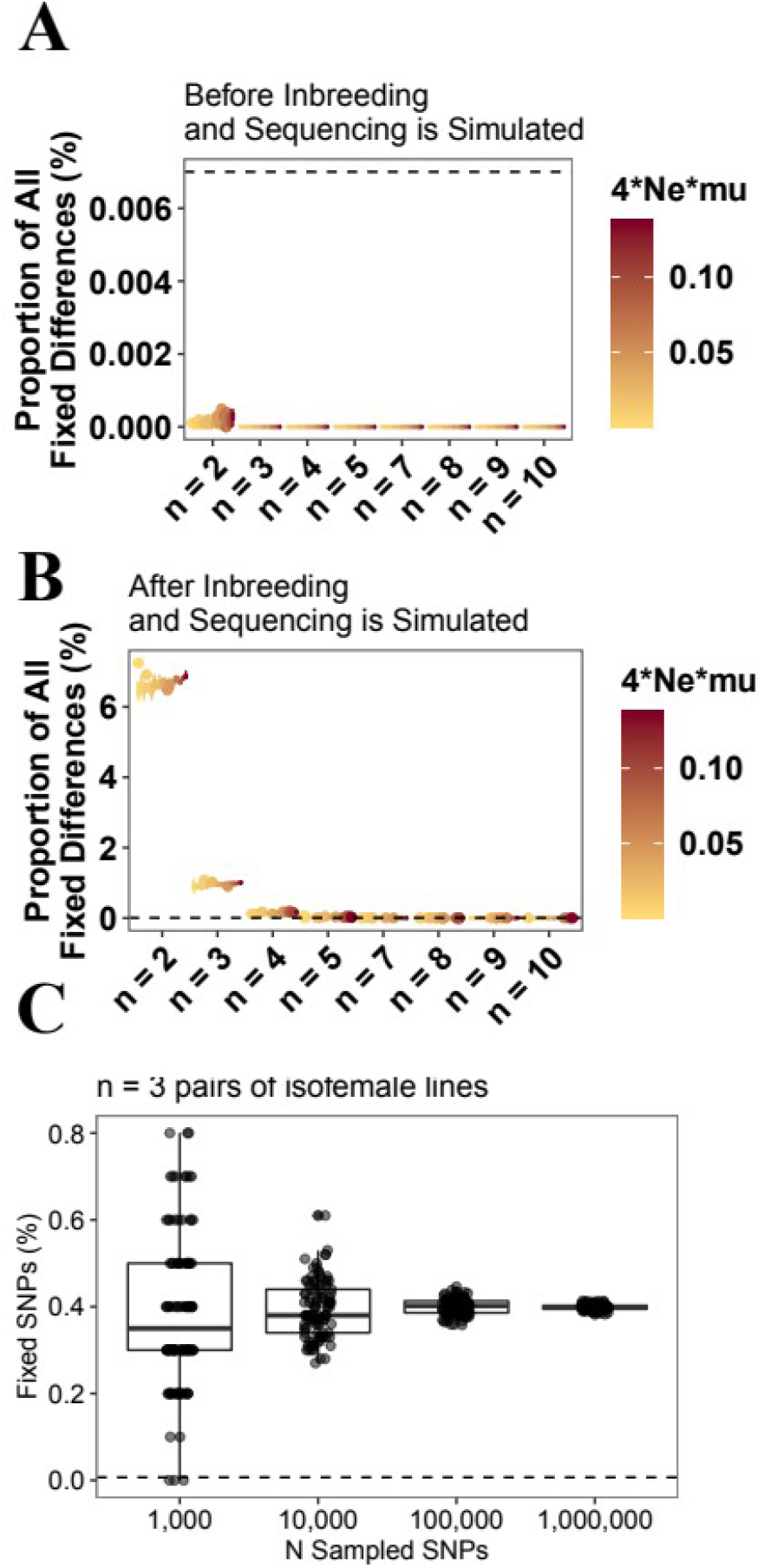
The proportion of all SNPs that are consistently fixed between *n* pairs of isofemale lines. Data are from population genetic simulations with different parameter combinations. **A** Results without simulating the inbreeding and lab maintenance design, **B** Results when this experimental design is taken into account. The *x*-axes in **A** and **B** show different numbers of isofemale lines pairs (*n*). Simulations were run for a range of values of *4Neμ.* **C** The expected proportion of all SNPs which are consistently fixed differences between *n* = 3 pairs of isofemale lines. The proportions are based on samples of 1,000, 10,000, 100,000, or 1,000,000 SNPs, variation in the estimates comes from 100 bootstraps. In all plots the horizontal dashed line gives the results seen in the pool-seq data (193 SNPs, 0.007%).

## Empirical Results

Although there is no association between chromosome length and the number of fixed SNPs (Spearman rank correlation: rho = 0.42, S = 126.6, p = 0.19) there is a strong correlation with the number of SNPs overall (rho = 0.74, S = 145.5, p = 0.0016). The genomic locations of the fixed SNPs are given in figure 3. The largest concentrations of fixed SNPs is on the 4^th^ chromosome (40% of all fixed SNPs), followed by both arms of the X chromosome (XR = 27%; XL = 10%). If the proportions of all SNPs on each chromosome is taken as the expected proportion of SNPs in a chi-squared test then there is a significant deviation from the expected distribution of top SNPs across chromosomes (Chi-squared = 145.76, d.f. = 14, p < 0.001) due to an excess on the 4^th^ and the X chromosomes.

**Figure 3.**
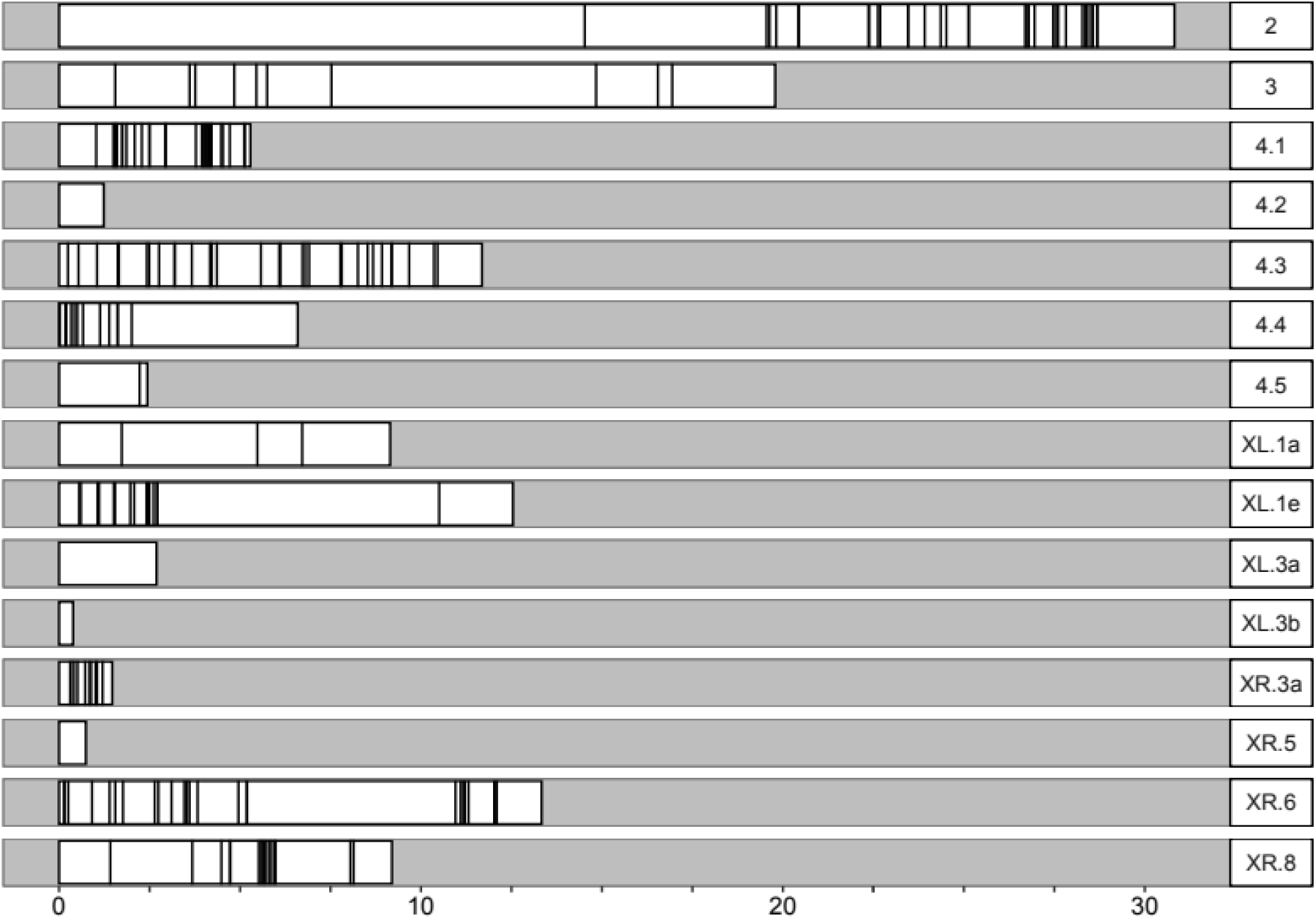
Chromosomal locations of 193 SNPs with consistently fixed differences between isofemale lines. Panel titles give the chromosome names. For chromosome 4 and the X chromosome arms the chromosomes are split into groups (see *Methods*’).

### Gene Ontology, Phenotype Enrichment, and TF Motif Enrichment

GO term enrichment analysis finds no enrichment of GO terms for genes within 1Mb of all fixed SNPs (all p > 0.05 after correction for multiple testing; table S3). In total 8,481 GO terms were tested for enrichment (table S3). The 193 fixed SNPs lie within 1Mb of 157 unique genes (table S4). In a phenotype enrichment analysis through modPhEA (Weng & Liao, 2017) genes within 1Mb of a fixed SNP are significantly enriched for various phenotypic classes (table S5). In total, 1,098 classes are tested and 65 pass a Benjamini-Hochberg correction for multiple testing (table S5). These include “behaviour defective” (FDR adjusted p-value = 0.008), and “female reproductive system” (p = 0.04). Of particular note are the genes within the “behaviour defective” class (table S6). One of these genes, *spineless (ss),* has mutant phenotypes which cause defective mating or courtship behaviours in *D. melanogaster*. No TF binding site motifs are enriched around fixed SNPs. A total of 656 motifs were tested and 178 motifs were significantly enriched (Fisher’s exact test, p < 0.05). However, none of these passed a correction for multiple testing (table S7). Nevertheless, motifs for *lola* are enriched (8 motifs with p-values < 0.05).

## DISCUSSION

Here we have used a method akin to replicated bulk segregant analysis to ask if we can find SNPs associated with a complex polygenic trait, female re-mating rate, from isofemale lines collected from the field. Although we only use three pairs of lines from three populations, and the lines were not necessarily the most extreme from the phenotypic distributions measured, we identify about 200 SNPs.

An important question is whether the ~200 SNPs identified differ from expectations given random sampling. We use both simulations and bootstrapping approaches to obtain estimates of the number of consistently fixed differences that are expected under a neutral model, with surprising results. We found many fewer fixed differences than population genetic simulations suggest, given an explicit model of the breeding history of the lines. Is there a reason the models should poorly predict the expected number of fixed differences? Females sampled from a population will have mated with at least one male whose genotype will also be represented among the offspring (David et al., 2005; Price et al., 2011). This should increase the genetic diversity within the isofemale line and reduce the probability of finding a fixed difference with another isofemale line by chance alone. If a female has mated with more than one male the genetic diversity among her offspring will be higher still and, consequently, the probability of finding a fixed difference with another isofemale line by chance is further reduced. On the other hand, inbreeding during laboratory maintenance will reduce diversity within the lines. Additionally, homozygous lethal alleles will never be fixed in isofemale lines thus some residual heterozygosity is always expected.

In the bootstrapping approach the distribution of allele frequencies from which bootstrap samples are drawn is a reflection of the variation among the isofemale lines in this study. Because these lines reflect only a subset of the variation found in wild populations the possible diversity is greatly reduced and our expected proportion of fixed differences is considerably less, though still substantially greater than the number actually observed.

A potential problem with both of the above approaches is that they do not account for linkage between SNPs. Closely linked SNPs will be in linkage disequilibrium (LD) and their allele frequencies highly correlated. Since many of the fixed SNPs in this study seem to occur in clusters (figure 3) this would inflate the number of fixed SNPs above the expected number from simulations, the opposite result from what we see. However, the fact that pairs of lines are from source populations all more than 500km apart should ensure that the different recombination histories has broken down LD, even between physically very close SNPs.

The poor performance of our simulations is unexpected. These are based on established population genetic equations describing the behaviour of alleles expected under neutral drift and mutation. In addition we parameterised these simulations with a range of mutation rates and effective population sizes that reflect the best estimates from natural populations of species similar to *D. pseudoobscura.* Variation in these parameters does not change the conclusions of this study. Therefore, we cannot conclude that we have identified more fixed SNPs than are expected by chance alone. Our population genetic simulations suggest that we would need at least 5 independent pairs of isofemale lines from different populations in order for the observed number of fixations (193) to be considered “significantly” more than expected if the models are accurate. Alternatively, biases in the bioinformatic pipeline could be to blame if decisions taken greatly increase the number of spurious SNPs called. We have attempted to follow best-practice guidelines (e.g. Schlötterer et al., 2014) but perhaps these guidelines need revisiting. Finally, if genetic diversity in these isofemale lines is much higher than expected from the inbreeding and lab maintenance design, this could explain the results.

### Genes and Regulatory Motifs Near Fixed SNPs

Another approach to asking if the study is successfully identifying candidate genes is to complete functional analyses of those found – do they implicate appropriate genes? The SNPs are spread throughout the chromosomes (figure 3) but are disproportionately found on chromosomes 4 & X. There is little evidence that they lie near or within genes previously implicated in mating behaviour phenotypes. The genes *pumilio* (*pum*) and *spineless* (*ss*) have previously been associated with variation in mating or courtship behaviour (Mackay et al., 2005). However, some other phenotypic terms are enriched among the top genes including “female reproductive system” and “behaviour defective” (table S5).

We also examined transcription factor (TF) binding site motifs and found that, after correcting for multiple testing, no TF motifs are enriched in regions around the fixed SNPs. However, several transcription factor motifs achieve marginal significance (p-values < 0.05 prior to correction for multiple testing. In particular, *lola* motifs are overrepresented among the transcription factor binding motifs. *lola* has been shown to change expression patterns in selection lines for slow and fast mating latency (Mackay et al., 2005).

Given the small number of lines involved, and the results of the simulation studies these results need to be treated with caution. Additional pairs of isofemale lines from different populations would be greatly beneficial. Additionally, female re-mating rate in *D. pseudoobscura* varies clinally (Price et al., 2014), and it would be very interesting to know if the SNPs identified here also show clinal variation. Clearly more lines, and pairwise comparisons, will further reduce the likelihood of false positives.

In conclusion, this study explores a method to identify SNPs associated with a complex quantitative trait, female re-mating rate. Isofemale lines from different populations, which differ in their re-mating rates, were pool-sequenced to identify markers consistently fixed for alternative alleles across the high and low re-mating lines. Population genetic simulations suggest that the number of fixed differences observed is consistently fewer than expected by chance. Further work is needed to demonstrate the feasibility of this relatively simple experimental breeding and selective sequencing approach to uncover loci associated with quantitative traits and to understand the poor fit of simulated data to the observed allele frequency differences.

## Supporting information

Analysis scripts and data

## Acknowledgments

This work was supported by a combined Natural Environment Research Council and St Andrews 600th Anniversary PhD Studentship grant (NE/L501852/1) to RAWW, a NERC grant (NE/J020818/1) to MGR, a NERC grant (NE/I027711/1) to NW, and a NERC fellowship (NE/H015684/1) to TARP. In addition the authors would like to thank Louise Reynolds, Paulina Giraldo-Perez, and Rudi Verspoor for help in the laboratory. Darren Obbard made helpful comments on a previous version of the manuscript, and recommended revisions to the simulations. Bioinformatics and Computational Biology analyses were supported by the University of St Andrews Bioinformatics Unit which is funded by a Wellcome Trust ISSF award (grant 105621/Z/14/Z).

## Conflict of Interest

The authors declare no conflicts of interest.

## Data Archiving

Raw reads have been deposited with NCBI under the BioProject accession: PRJNA602380. Analysis scripts and simulation data are included in an archive as a supplement to this manuscript.

## Notes

### Competing Interest Statement

The authors have declared no competing interest.

## REFERENCES

Bailey TL, Boden M, Buske FA, Frith M, Grant CE, Clementi L, et al (2009). MEME Suite: Tools for motif discovery and searching. Nucleic Acids Res 37: 202–208.

Bergland AO, Behrman EL, O’Brien KR, Schmidt PS, Petrov DA (2014). Genomic evidence of rapid and stable adaptive oscillations over seasonal time scales in *Drosophila*. PLoS Genet 10: e1004775.

Berriz GF, Beaver JE, Cenik C, Tasan M, Roth FP (2009). Next generation software for functional trend analysis. Bioinformatics 25: 3043–3044.

Bhattacharjee MJ, Yu C-P, Lin J-J, Ng CS, Wang T-Y, Lin H-H, Li W-H (2016). Regulatory divergence among beta-keratin genes during bird evolution. Mol Biol Evol 33: 2769–2780.

Boake CRB, Arnold SJ, Breden F, Meffert LM, Ritchie MG, et al (2002). Genetic tools for studying adaptation and the evolution of behavior. Am Nat 160: S143–59.

Bolger AM, Lohse M, Usadel B (2014). Trimmomatic: a flexible trimmer for Illumina sequence data. Bioinformatics 30: 2114–2120.

Broad Institute Picard v. 2.14.1. url: http://broadinstitute.github.io/picard/

Burke MK, Dunham JP, Shahrestani P, Thornton KR, Rose MR, Long AD (2010). Genome wide analysis of a long-term evolution experiment with *Drosophila*. Nature 467: 587–90.

Chan YF, Marks ME, Jones FC, Villareal Jr, G, Shapiro MD, Brady SD, et al (2010). Adaptive evolution of pelvic reduction in sticklebacks by recurrent deletion of a *Pitx1* enhancer. Science 327: 302–306.

Chapman T, Bangham J, Vinti G, Seifried B, Lung O, Wolfner MF, et al (2003). The sex peptide of *Drosophila melanogaster*: Female post-mating responses analyzed by using RNA interference. Proc Natl Acad Sci U S A 100: 9923–9928.

Charlesworth B, Charlesworth D (2008). Elements of Evolutionary Genetics. Roberts and Company Publishers, Greenwood Village, Colorado.

Chen J, Källman T, Ma X-F, Zain G, Morgante M, Lascoux M (2016). Identifying genetic signatures of natural selection using pooled population sequencing in *Picea abies*. G3 6: 1979–1989.

Colosimo PF, Péichel CL, Nereng K, Blackman BK, Shapiro MD, Schluter D, Kingsley DM (2004). The genetic architecture of parallel armor plate reduction in threespine sticklebacks. PLoS Biol 2: e0635.

David JR, Gibert P, Legout H, Pétavy G, Capy P, Moreteau B (2005). Isofemale lines in *Drosophila*: an empirical approach to quantitative trait analysis in natural populations. Heredity. 94: 3–12.

DePristo MA, Banks E, Poplin R, Garimella KV, Maguire JR, Hartl C, et al (2011). A framework for variation discovery and genotyping using next-generation DNA sequencing data. Nat Genet 43: 491–498.

dos Santos G, Schroeder AJ, Goodman JL, Strelets VB, Crosby MA, Thurmond J, et al (2014). FlyBase: introduction of the *Drosophila melanogaster* Release 6 reference genome assembly and large-scale migration of genome annotations. Nucleic Acids Res 43: D690–D697.

Giardina TJ (2015). Mating rate and the influence of female genetics on remating in *Drosophila melanogaster*. PhD Thesis. Binghamton University.

Giardina TJ, Beavis A, Clark AG, Fiumera AC (2011). Female influence on pre-and postcopulatory sexual selection and its genetic basis in *Drosophila melanogaster*. Mol Ecol 20: 4098–4108.

Haag-Liautard C, Dorris M, Maside X, Macaskill S, Halligan DL, Houle D, et al (2007). Direct estimation of per nucleotide and genomic deleterious mutation rates in *Drosophila*. Nature 445: 82–85.

Haerty W, Jagadeeshan S, Kulathinal RJ, Wong A, Ravi Ram K, Sirot LK, et al (2007). Evolution in the fast lane: rapidly evolving sex-related genes in *Drosophila*. Genetics 177: 1321–1335.

Hoban S, Kelley JL, Lotterhos KE, Antolin MF, Bradburd G, Lowry DB, et al (2016). Finding the genomic basis of local adaptation: pitfalls, practical solutions, and future directions. Am Nat 188: 379–397.

Huang W, Massouras A, Inoue Y, Peiffer J, Ràmia M, Tarone A, et al (2014). Natural variation in genome architecture among 205 *Drosophila melanogaster* Genetic Reference Panel lines. Genome Res 24: 1193–1208.

Ivanov DK, Escott-Price V, Ziehm M, Magwire MM, Mackay TFC, Partridge L, Thornton JM (2015). Longevity GWAS using the *Drosophila* Genetic Reference Panel. J Gerontol A Biol Sci Med Sci 70: 1470–1478.

Jensen JD, Bachtrog D (2011). Characterizing the Influence of effective population size on the rate of adaptation: Gillespie’s Darwin Domain. Genome Biol and Evol 3: 687–701.

Keightley PD, Ness RW, Halligan DL, Haddrill PR (2014). Estimation of the spontaneous mutation rate per nucleotide site in a *Drosophila melanogaster* full-sib family. Genetics 196: 313–320.

Keightley PD, Pinharanda A, Ness RW, Simpson F, Dasmahapatra KK, Mallet J, et al (2015). Estimation of the spontaneous mutation rate in *Heliconius melpomene*. Mol Biol Evol 32: 239–243.

Kleinjan DA, Heyningen V Van. (2005). Long-Range Control of Gene Expression: Emerging mechanisms and disruption in disease. Am J Hum Genet 76: 8–32.

Kofler R, Pandey RV, Schlötterer C (2011). PoPoolation2: identifying differentiation between populations using sequencing of pooled DNA samples (Pool-Seq). Bioinformatics 27: 3435–3436.

Kofler R, Schlötterer C (2014). A guide for the design of evolve and resequencing studies. Mol Biol Evol 31: 474–483.

Kofler R, Schlötterer C (2012). Gowinda: Unbiased analysis of gene set enrichment for genome-wide association studies. Bioinformatics 28: 2084–2085.

Kronforst MR, Young LG, Kapan DD, Mcneely C, Neill RJO, Gilbert LE (2006). Linkage of butterfly mate preference and wing color preference cue at the genomic location of wingless. Proc Natl Acad Sci U S A 103: 6575–6580.

Lamichhaney S, Martinez Barrio A, Rafati N, Sundström G, Rubin C-J, Gilbert ER, et al (2012). Population-scale sequencing reveals genetic differentiation due to local adaptation in Atlantic herring. Proc Natl Acad Sci U S A109: 19345–50.

Leal WS (2013). Odorant Reception in Insects: Roles of receptors, binding proteins, and degrading enzymes. Annu Rev Entomol 58.

Li H (2013). Aligning sequence reads clone sequences and assembly contigs with BWA-MEM *arXiv Prepr*. arXiv 0: 3.

Li H, Handsaker B, Wysoker A, Fennell T, Ruan J, Homer N, et al (2009). The Sequence Alignment/Map format and SAMtools. Bioinformatics 25: 2078–2079.

Mackay TFC, Heinsohn SL, Lyman RF, Amanda J, Morgan TJ, Rollmann SM, et al (2005). Genetics and genomics of *Drosophila* mating behavior. Proc Natl Acad Sci U S A 102: 6622–6629.

Mackay TFC, Richards S, Stone EA, Barbadilla A, Ayroles JF, Zhu D, et al (2012). The *Drosophila melanogaster* Genetic Reference Panel. Nature 482: 173–178.

Magwene PM, Willis JH, Kelly JK (2011). The statistics of bulk segregant analysis using next generation sequencing. PLoS Comput Biol 7: e1002255.

Maston GA, Evans SK, Green MR (2006). Transcriptional regulatory elements in the human genome. Annu Rev Genomics Hum Genet 7: 29–59.

McGraw LA, Gibson G, Clark AG, Wolfner, MF (2004). Genes regulated by mating, sperm, or seminal proteins in mated female *Drosophila melanogaster*. Currr Biol 14: 1509–1514.

McGraw L a., Clark AG, Wolfner MF (2008). Post-mating gene expression profiles of female *Drosophila melanogaster* in response to time and to four male accessory gland proteins. Genetics 179: 1395–1408.

McKenna A, Hanna M, Banks E, Sivachenko A, Cibulskis K, Kernytsky A, et al (2010). The Genome Analysis Toolkit: A MapReduce framework for analyzing next-generation DNA sequencing data. Genome Res 20: 1297–1303.

McLeay RC, Bailey TL (2010). Motif Enrichment Analysis: a unified framework and an evaluation on ChIP data. BMC Bioinformatics. 11: 165.

Montgomery SL, Vorojeikina D, Huang W, Mackay TFC, Anholt RRH, Rand MD (2014). Genome-wide association analysis of tolerance to methylmercury toxicity in *Drosophila* implicates myogenic and neuromuscular developmental pathways. PLoS ONE 9: e110375

Neville MC, Nojima T, Ashley E, Parker DJ, Walker J, Southall T, et al (2014). Male-specific fruitless isoforms target neurodevelopmental genes to specify a sexually dimorphic nervous system. Curr Biol 24: 229–241.

Noor MAF, Schug MD, Aquadro CF (2000). Microsatellite variation in populations of *Drosophila pseudoobscura* and *Drosophila persimilis*. Genetics Res 75: 25–35.

Orozco-terWengel P, Kapun M, Nolte V, Kofler R, Flatt T, Schlötterer C (2012). Adaptation of *Drosophila* to a novel laboratory environment reveals temporally heterogeneous trajectories of selected alleles. Mol Ecol 21: 4931–4941.

Pennacchio LA, Bickmore W, Dean A, Nobrega MA, Bejerano G (2013). Enhancers: five essential questions. Nat Rev Genet 14: 288–295.

Pizzari T, Wedell N (2013). The polyandry revolution. Philos Trans R Soc Lond B Biol Sci 368: 20120041.

Price TAR, Bretman A, Gradilla AC, Reger J, Taylor ML, Campbell A, et al (2014). Does polyandry control population sex ratio via regulation of a selfish gene? Proc Biol Sci 281: 20133259.

Price TAR, Hodgson DJ, Lewis Z, Hurst GDD, Wedell N (2008). Selfish genetic elements promote polyandry in a fly. Science. 211: 1241–1243.

Price TAR, Hurst GDD, Wedell N (2010). Polyandry prevents extinction. Curr Biol. 20: 471–475.

Price TAR, Lewis Z, Smith DT, Hurst GDD, Wedell N (2010). Sex ratio drive promotes sexual conflict and sexual coevolution in the fly *Drosophila pseudoobscura*. Evolution. 64: 1504–1509.

Price TAR, Lewis Z, Smith DT, Hurst GDD, Wedell N (2011). Remating in the laboratory reflects rates of polyandry in the wild. Animal Behav 82: 1381–1386.

Quinlan AR, Hall IM (2010). BEDTools: a flexible suite of utilities for comparing genomic features. Bioinformatics 26: 841–842.

Ram KR, Wolfner MF (2007). Seminal influences: *Drosophila* Acps and the molecular interplay between males and females during reproduction. Integr Comp Biol 47: 427–445.

Rockman M V (2011). The QTN program and the alleles that matter for evolution: all that’s gold does not glitter. Evolution. 66: 1–17.

Schlötterer C, Kofler R, Versace E, Tobler R, Franssen SU (2015). Combining experimental evolution with next-generation sequencing: a powerful tool to study adaptation from standing genetic variation. Heredity. 114: 431–440.

Schlötterer C, Tobler R, Kofler R, Nolte V (2014). Sequencing pools of individuals — mining genome-wide polymorphism data without big funding. Nat Rev Genet 15: 749–763.

Schneeberger K (2014). Using next-generation sequencing to isolate mutant genes from forward genetic screens. Nat Rev Genet 15: 662–676.

Snook RR (2014). The evolution of polyandry. In: DM Shuker & LW Simmons (eds) The Evolution of Insect Mating Systems. Oxford University Press, Oxford, Pp 159–180

Sokolowski MB (2001). *Drosophila*: genetics meets behaviour. Nat Rev Genet 2: 879–890.

Stapley J, Reger J, Feulner PGD, Smadja C, Galindo J, Ekblom R, et al (2010). Adaptation genomics: the next generation. Trends Ecol Evol 25: 705–712.

Swanson WJ, Wong A, Wolfner MF, Aquadro CF (2004). Evolutionary expressed sequence tag analysis of *Drosophila* female reproductive tracts identifies genes subjected to positive selection. Genetics 168: 1457–1465.

The 1001 Genomes Consortium (2016). 1,135 Genomes Reveal the Global Pattern of Polymorphism in *Arabidopsis thaliana*. Cell 166: 481–491.

Tram U, Wolfner MF (1998). Seminal fluid regulation of female sexual attractiveness in *Drosophila melanogaster*. Proc Natl Acad Sci U S A 95: 4051–4054.

Travisano M, Shaw RG (2012). Lost in the map. Evolution. 67: 305–314.

Tsuda M, Aigaki T (2016). Evolution of sex-peptide in *Drosophila*. Fly. 10: 172–177.

Tsuda M, Peyre JB, Asano T, Aigaki T (2015). Visualizing molecular functions and crossspecies activity of sex-peptide in *Drosophila*. Genetics 200: 1161–1169.

Weng MP, Liao BY (2017). modPhEA: model organism Phenotype Enrichment Analysis of Eukaryotic gene sets. Bioinformatics 33: 3505–3507.

Werner T, Koshikawa S, Williams TM, Carroll SB (2010). Generation of a novel wing colour pattern by the *Wingless* morphogen. Nature 464: 1143–1148.

Wittkopp PJ, Kalay G (2012). Cis-regulatory elements: molecular mechanisms and evolutionary processes underlying divergence. Nat Rev Genet 13: 59–69.

Yang J, Jiang H, Yeh CT, Yu J, Jeddeloh JA, Nettleton D, et al (2015). Extreme-phenotype genome-wide association study (XP-GWAS): A method for identifying trait-associated variants by sequencing pools of individuals selected from a diversity panel. Plant J 84: 587–596.

Yapici N, Kim Y-J, Ribeiro C, Dickson BJ (2008). A receptor that mediates the post-mating switch in *Drosophila* reproductive behaviour. Nature 451: 33–37.

