## Supplementary material for "Identifying genomic markers associated with female re-mating rate in *Drosophila pseudoobscura* by replicated bulk segregant analysis": Analysis scripts and data: pseudo_wild_alfreq_analysis.R

```
##
# D.pseudoobscura project wild linse
# PoPoolation2 allele frequencies data analysis
# Last Modified: March 2017
##
#clean environment
rm(list = ls(all = TRUE))
#

# Load libraries and define functions
# #####
library(plyr)
library(ggplot2)
library(scales)
#install.packages("grid")
library(gridExtra)
library(grid)
#install.packages("reshape")
library(reshape)
library(dplyr)
se <- function(x) {sqrt(var(x, na.rm = TRUE))/sqrt(length(x))}
setwd("~/Dropbox/st_andrews/pseudoobscura_wild_lines_MS/BioRxiv_submission/scripts_data/")
source("~/projects/RData/RScripts/ggplot_theme.R")
options(scipen = 6)

# Read in data
# Load data
al_freq_dat <- read.table(
  "~/Dropbox/st_andrews/pseudoobscura_wild_lines_MS/BioRxiv_submission/scripts_data/pseudo_wild_SHA-SLOB7subs_filt_alfreq.tab",
                          sep = "\t", header = FALSE)
colnames(al_freq_dat) <- c("Chr","Pos",
                           "ref","maj",
                           "2SLOC9","SLOB7",
                           "LEW17","LEW23",
                           "SHAC1","SHAA10")
head(al_freq_dat)
al_freq_dat_m<-melt(al_freq_dat[,c(5,6,7,8,9)])
head(al_freq_dat_m)
high<-c("LEW23","SHAC1","SLOB7")
low<-c("LEW17","SHAA10","2SLOC9")
al_freq_dat<-cbind(al_freq_dat,al_freq_dat[,high]-al_freq_dat[,low])
colnames(al_freq_dat)<-c("Chr","Pos",
                         "ref","maj",
                         "2SLOC9","SLOB7",
                         "LEW17","LEW23",
                         "SHAC1","SHAA10","Lew","Sha","Slo")
nrow(al_freq_dat)
# Subset to those on known chromosomes
al_freq_dat<-al_freq_dat[grep("Unknown",al_freq_dat$Chr,invert=TRUE),]
nrow(al_freq_dat)

nrow(al_freq_dat[which(al_freq_dat$Slo == 1 & 
                         al_freq_dat$Lew == 1 & 
                         al_freq_dat$Sha == 1),])+
nrow(al_freq_dat[which(al_freq_dat$Slo == -1 & 
                         al_freq_dat$Lew == -1 & 
                         al_freq_dat$Sha == -1),])

n_fixed_diffs<-nrow(al_freq_dat[which(al_freq_dat$Slo == 1 & 
                         al_freq_dat$Lew == 1 & 
                         al_freq_dat$Sha == 1),])+
  nrow(al_freq_dat[which(al_freq_dat$Slo == -1 & 
                           al_freq_dat$Lew == -1 & 
                           al_freq_dat$Sha == -1),])
nrow(al_freq_dat)

# As a percentage of all SNPs
(n_fixed_diffs/nrow(al_freq_dat))*100

al_freq_dat$Chr2<-gsub("chrom","",al_freq_dat$Chr)
al_freq_dat$Chr2<-gsub("_group",".",al_freq_dat$Chr2)
al_freq_dat$Chr3<-gsub("\\..*","",al_freq_dat$Chr2)

al_freq_dat$snp<-paste("SNP_",seq(1,nrow(al_freq_dat)),sep="")
# Subset to those SNPs that are fixed in all populations
alfreq_fixed<-rbind(al_freq_dat[which(al_freq_dat$Slo == 1 & 
                                  al_freq_dat$Lew == 1 & 
                                  al_freq_dat$Sha == 1),],
                    al_freq_dat[which(al_freq_dat$Slo == -1 & 
                                        al_freq_dat$Lew == -1 & 
                                        al_freq_dat$Sha == -1),])
                    

nrow(al_freq_dat)
(nrow(alfreq_fixed)/nrow(al_freq_dat))*100

tapply(alfreq_fixed$Chr,INDEX = list(alfreq_fixed$Chr3),length)

tapply(alfreq_fixed$Chr,INDEX = list(alfreq_fixed$Chr3),length)/nrow(alfreq_fixed)*100

# Plot the fixed SNPs

# Chromosome lengths
main_chroms <- read.table(
  file = "~/projects/pseudo_reference/pseudo_main_chrom_data.tab")


colnames(main_chroms)<-c("Chr","type","coords","name","sum","length","version")
main_chroms$Chr2<-gsub("_group",".",main_chroms$Chr)
head(main_chroms)

# Correlate chrom length to nr snps
nr_snps<-tapply(al_freq_dat$Chr,INDEX = list(al_freq_dat$Chr2),length)

nr_fixed_snps<-tapply(alfreq_fixed$Chr,INDEX = list(alfreq_fixed$Chr2),length)
nr_fixed_snps<-nr_fixed_snps[match(names(nr_snps),names(nr_fixed_snps))]
names(nr_fixed_snps)<-names(nr_snps)
nr_fixed_snps[which(is.na(nr_fixed_snps))]<-0

main_chroms$length[match(names(nr_snps),main_chroms$Chr2)]

cor.test(x=nr_fixed_snps,y=main_chroms$length[match(names(nr_snps),main_chroms$Chr2)],method = "spearman")
cor.test(x=nr_fixed_snps,y=nr_snps,method = "spearman")


# Chi squared test.


p<-nr_snps/nrow(al_freq_dat)

t<-chisq.test(nr_fixed_snps,p=p)
t$observed[t$observed>t$expected]

# Make an ideogrammatic chromosome plot with fixed SNPs
ggplot()+
  geom_rect(data=main_chroms,aes(xmin=0,xmax=length/1000000,ymin=0,ymax=1),
            fill="white",colour="black")+
  geom_vline(data=alfreq_fixed,aes(xintercept = Pos/1000000))+
  scale_x_continuous(breaks=c(0,2.5,5,7.5,10,12.5,15,17.5,20,22.5,25,27.5,30,22.5),
                     labels=c("0","","","","10","","","","20","","","","30",""))+
  ylim(0,1)+
  facet_grid(Chr2~.)+
  my.theme+
  theme(panel.background = element_rect(fill="grey"),
        panel.grid.major=element_blank(),
        panel.grid.minor=element_blank(),
        axis.text.y = element_blank(),
        axis.title.y = element_blank(),
        axis.ticks.y = element_blank(),
        strip.text.y = element_text(angle = 0,size = 10))

tapply(alfreq_fixed$Chr,INDEX = list(alfreq_fixed$Chr3),length)

# How many sites show at least one fixation
nrow(alfreq[alfreq$SLo == 1 | 
              alfreq$Lew == 1 | 
              alfreq$Sha == 1,])
##------------------------------------------------------------##
# WRITE FIXED SNPS TO GTF FILE
##------------------------------------------------------------##
head(al_freq_dat)
nrow(al_freq_dat)

head(alfreq_fixed)
nrow(alfreq_fixed)

gtf<-data.frame("chr"=al_freq_dat$Chr,
                "source"=rep("PoPoolation2",nrow(al_freq_dat)),
                "type"=rep("CDS",nrow(al_freq_dat)),
                "start"=al_freq_dat$Pos,
                "end"=al_freq_dat$Pos,
                "score"=rep(".",nrow(al_freq_dat)),
                "strand"=rep(".",nrow(al_freq_dat)),
                "X"=rep(".",nrow(al_freq_dat)),
                "feature"=paste('gene_id "',al_freq_dat$snp,
                                '"; transcript_id "',al_freq_dat$snp,'"',sep=""))
head(gtf)
write.table(gtf,"~/projects/pseudo_wild/SNPs_analysis/pseudo_wild_all_snps.gtf",
            col.names=FALSE,row.names=FALSE,quote=FALSE,sep="\t")

fixed_gtf<-data.frame("chr"=alfreq_fixed$Chr,
                "source"=rep("PoPoolation2",nrow(alfreq_fixed)),
                "type"=rep("CDS",nrow(alfreq_fixed)),
                "start"=alfreq_fixed$Pos,
                "end"=alfreq_fixed$Pos,
                "score"=rep(".",nrow(alfreq_fixed)),
                "strand"=rep(".",nrow(alfreq_fixed)),
                "X"=rep(".",nrow(alfreq_fixed)),
                "feature"=paste('gene_id "',alfreq_fixed$snp,
                                '"; transcript_id "',alfreq_fixed$snp,'"',sep=""))
head(fixed_gtf)
write.table(fixed_gtf,"~/projects/pseudo_wild/SNPs_analysis/pseudo_wild_fixed_snps.gtf",
            col.names=FALSE,row.names=FALSE,quote=FALSE,sep="\t")

options(scipen = 10)
gtf<-data.frame("chr"=al_freq_dat$Chr,
                "source"=rep("PoPoolation2",nrow(al_freq_dat)),
                "type"=rep("CDS",nrow(al_freq_dat)),
                "start"=al_freq_dat$Pos-30,
                "end"=al_freq_dat$Pos+30,
                "score"=rep(".",nrow(al_freq_dat)),
                "strand"=rep(".",nrow(al_freq_dat)),
                "X"=rep(".",nrow(al_freq_dat)),
                "feature"=paste('gene_id "',al_freq_dat$snp,
                                '"; transcript_id "',al_freq_dat$snp,'"',sep=""))
head(gtf)
write.table(gtf,"~/projects/pseudo_wild/SNPs_analysis/pseudo_wild_all_snps_30bp.gtf",
            col.names=FALSE,row.names=FALSE,quote=FALSE,sep="\t")

fixed_gtf<-data.frame("chr"=alfreq_fixed$Chr,
                      "source"=rep("PoPoolation2",nrow(alfreq_fixed)),
                      "type"=rep("CDS",nrow(alfreq_fixed)),
                      "start"=alfreq_fixed$Pos-30,
                      "end"=alfreq_fixed$Pos+30,
                      "score"=rep(".",nrow(alfreq_fixed)),
                      "strand"=rep(".",nrow(alfreq_fixed)),
                      "X"=rep(".",nrow(alfreq_fixed)),
                      "feature"=paste('gene_id "',alfreq_fixed$snp,
                                      '"; transcript_id "',alfreq_fixed$snp,'"',sep=""))
head(fixed_gtf)
write.table(fixed_gtf,"~/projects/pseudo_wild/SNPs_analysis/pseudo_wild_fixed_snps_30bp.gtf",
            col.names=FALSE,row.names=FALSE,quote=FALSE,sep="\t")
X<-2000
fixed_gtf<-data.frame("chr"=alfreq_fixed$Chr,
                      "source"=rep("PoPoolation2",nrow(alfreq_fixed)),
                      "type"=rep("CDS",nrow(alfreq_fixed)),
                      "start"=alfreq_fixed$Pos-X,
                      "end"=alfreq_fixed$Pos+X,
                      "score"=rep(".",nrow(alfreq_fixed)),
                      "strand"=rep(".",nrow(alfreq_fixed)),
                      "X"=rep(".",nrow(alfreq_fixed)),
                      "feature"=paste('gene_id "',alfreq_fixed$snp,
                                      '"; transcript_id "',alfreq_fixed$snp,'"',sep=""))
head(fixed_gtf)

head(main_chroms)
fixed_gtf$start[which(fixed_gtf$start<0)]<-1
for(chr in main_chroms$Chr ){
  max_l<-main_chroms$length[which(main_chroms$Chr==chr)]
  fixed_gtf$end[which(fixed_gtf$chr == chr & fixed_gtf$end > max_l)]<-max_l
}

write.table(fixed_gtf,paste(
  "~/projects/pseudo_wild/SNPs_analysis/pseudo_wild_fixed_snps_",X,"bp.gtf",sep=""),
            col.names=FALSE,row.names=FALSE,quote=FALSE,sep="\t")

##------------------------------------------------------------##
# ALLELE FREQUENCY DISTRIBUTIONS IN EACH LINE
##------------------------------------------------------------##
# Load data
head(al_freq_dat)

al_freq_dat<-al_freq_dat[grep("Unknown",al_freq_dat$Chr,invert=TRUE),]

nrow(al_freq_dat[which(al_freq_dat$Slo == 1 & 
                    al_freq_dat$Lew == 1 & 
                    al_freq_dat$Sha == 1),])
nrow(al_freq_dat[which(al_freq_dat$Slo == -1 & 
                         al_freq_dat$Lew == -1 & 
                         al_freq_dat$Sha == -1),])
nrow(al_freq_dat)


nrow(al_freq_dat[which(abs(al_freq_dat$Slo) == 1 & 
                         abs(al_freq_dat$Lew) == 1 & 
                         abs(al_freq_dat$Sha) == 1),])

# Collect counts for allele frequencies in 20 even bins
pop_freq_counts<-data.frame()
for(pop in c("2SLOC9","SLOB7","LEW17","LEW23","SHAC1","SHAA10")){
  bins<-seq(0,1,0.05)
  hist<-hist(al_freq_dat[,pop],breaks = bins)
  hist$mids
  counts<-hist$counts
  freq_counts<-data.frame("bin"=hist$mids,
                          "count"=counts,
                          "pop"=rep(pop,length(bins[-1])))
  pop_freq_counts<-rbind(pop_freq_counts,freq_counts)
}
head(pop_freq_counts)

#Compute the aggregate sum
aggr_count<-aggregate(pop_freq_counts$count,by=list(pop_freq_counts$bin),sum)
pop_freq_counts<-rbind(pop_freq_counts,data.frame("bin"=aggr_count[,1],
                                                  "count"=aggr_count[,2],
                                                  "pop"=rep("Aggregate",nrow(aggr_count))))
pop_freq_counts$pop<-factor(pop_freq_counts$pop,
                            levels = c("LEW17","LEW23","SHAA10","SHAC1",
                                       "SLOB7","2SLOC9","Aggregate"),
                            labels=c("Lewistown\n(Low; LEW17)",
                                     "Lewistown\n(High; LEW23)",
                                     "Shaver Lake\n(Low; SHAA10)",
                                     "Shaver Lake\n(High; SHAC1)",
                                     "Show Low\n(Low; 2SLOC9)",
                                     "Show Low\n(High; SLOB7)",
                                     "Aggregate"))
ggplot()+
  geom_bar(data=pop_freq_counts,
                 aes(x=bin,y=count/1000000),stat="identity")+
  facet_grid(pop~.)+
  ylab(expression(paste("Count (x10"^6,")",sep="")))+
  xlab("Allele Frequency")+
  my.theme+
  theme(axis.text.x = element_text(size=12,face="bold"),
        axis.text.y = element_text(size = 12,face = "bold"),
        strip.text.y = element_text(size = 12,angle = 0),
        #axis.ticks.x = element_blank(),
        legend.position = "top",
        legend.text = element_text(size=10,face="bold"),
        legend.title = element_text(size=10,face="bold"))

# Bootstrap sample from the aggregate allele frequency distribution to get expected allele frequency differences
aggr_count$prop<-aggr_count$x/sum(aggr_count)

n_lines<-3
n_snps<-c(1000,10000,100000,1000000)
n_sims<-100

# Define function for the sequencing.
sequencing<-function(x,draws){
  # Pooled sequencing (to target coverage 40x)
  # x is now the frequency in the tube at the time of sequencing
  # read-depth is poisson, with a mean of 50x
  rpois(draws,50)->depth
  rbinom(draws,depth,x)/depth->allele_frequency_in_reads
  #return the frequency after the experiment and sequencing
  return(allele_frequency_in_reads)
}
boot_dat<-data.frame("N_fixed"=vector(length=n_sims*length(n_snps)),
                     "N_snps"=vector(length=n_sims*length(n_snps)),
                     "boot"=vector(length=n_sims*length(n_snps)))
i<-1
for(n_snp in n_snps){
  sim<-1
  while(sim<=n_sims){
    high_ps<-vector(length=n_lines)
    high_gs<-vector(length=n_lines)
    low_ps<-vector(length=n_lines)
    low_gs<-vector(length=n_lines)
    #l=1
    #n_snp<-1000
    snp_highs<-matrix(nrow=n_snp,ncol = n_lines)
    snp_lows<-matrix(nrow=n_snp,ncol = n_lines)
    cat("n_snp: ", n_snp," - n_sims: ",sim,"\n")
    snp=1
    while(snp <= n_snp){
      for(l in 1:n_lines){
        # Sample allele frequencies for the 3 "high" lines
        high_ps<-sample(aggr_count[,1],size = 1,prob = aggr_count[,2])
        # Sample allele frequencies for the 3 "low" lines
        low_ps<-sample(aggr_count[,1],size = 1,prob = aggr_count[,2])
        # Obtain genotypes for the 3 "high" lines
        snp_highs[snp,l]<-mean(rbinom(n = 2,prob = high_ps,size = 1))
        # Obtain genotypes for the 3 "low" lines
        snp_lows[snp,l]<-mean(rbinom(n = 2,prob = low_ps,size = 1))
      }
      snp<-snp+1
    }
    boot_dat$N_fixed[i]<-length(which(apply(snp_highs-snp_lows,MARGIN = 1,
                                     function(x){all(x==1) | all(x==-1)})))
    boot_dat$N_snps[i]<-n_snp
    boot_dat$boot[i]<-sim
    sim<-sim+1
    i<-i+1
  }
}
write.table(boot_dat,"pseudo_wild_boots_p_dat.csv",
            col.names = TRUE,row.names=FALSE,quote=FALSE,sep=",")

head(boot_dat)
tapply((boot_dat$N_fixed/boot_dat$N_snps)*100,INDEX=list(as.character(boot_dat$N_snps)),quantile,0.95)
tapply((boot_dat$N_fixed/boot_dat$N_snps)*100,INDEX=list(as.character(boot_dat$N_snps)),quantile,0.05)

ggplot(data=boot_dat)+
  geom_boxplot(aes(as.character(N_snps),(N_fixed/N_snps)*100))+
  geom_point(aes(as.character(N_snps),(N_fixed/N_snps)*100),position=position_jitter(0.1),alpha=1/2)+
  xlab("N Sampled SNPs")+
  ggtitle("n = 3 pairs of isofemale lines")+
  scale_x_discrete("N Sampled SNPs",breaks=c(1000,10000,100000,1000000),
                   labels=c("1,000","10,000","100,000","1,000,000"))+
  ylab("Fixed SNPs (%)")+
  geom_hline(yintercept = 0.007,linetype="dashed",size=0.5)+
  my.theme


##------------------------------------------------------------##
# CLOSEST GENES
##------------------------------------------------------------##

snps_genes<-read.table(
  "~/Dropbox/st_andrews/pseudoobscura_wild_lines_MS/BioRxiv_submission/TableS4_fixed_snps_closest_genes_reduced.tab",
  sep="\t",
  header=FALSE)

colnames(snps_genes)<-c("chr","start","end","snp","score","strand","source","type","X","gene","distance")
head(snps_genes)
nrow(snps_genes)
length(unique(snps_genes$gene[which(abs(snps_genes$distance) < 1000000)]))
nrow(snps_genes[which(abs(snps_genes$distance) < 10000),])
```
