## Supplementary material for "Identifying genomic markers associated with female re-mating rate in *Drosophila pseudoobscura* by replicated bulk segregant analysis": Analysis scripts and data: pseudo_wild_alfreq_simulation_mod.R

```
# library(ggplot2)
# source("~/projects/RData/RScripts/ggplot_theme.R")
##------------------------------------------------------------##
# SIMULATIONS
# What proportion of sites do we expect to be fixed by chance
##------------------------------------------------------------##

# Describe the population
### H. melpomene:    ##
# Keightley et al., 2015 Molecular Biology and Evolution 32:239-243; Ne = 2*10^6
### D. melanogaster: ##
# Keightley et al., 2014 Genetics 196:313-320; Ne = 1.4*10^6
### D. pseudoobscura:##
# Schaeffer 1995 Genetics of Natural Populations: The Continuing Importance of Theodosius Dobzhansky; Ne = ??? (NOT AVAILABLE TO ME)
# Noor et al., 2000 Genetic Research 75:25-35; Ne = 141,000-512,000 (microsats)
# Jensen & Bachtrog 2011 Genome Biology and Evolution 3:687-701; Ne = 4.5*10^6 (nucleotide variation at 123 coding sequence fragments)
### Ne IS DEFINED IN THE SIMULATION: RUN FOR Ne = 1,2,3,and 4*10^6

### D. melanogaster: ##
# Haag-Liautard et al., 2007 Nature 445:82-85; u = 8.4*10^-9
# Keightley et al., 2014 Genetics 196:313-320; u = 2.8*10^-9 (95% CI = 1.0*10^-9 6.1*10^-9)
### H. melpomene: ##
# Keightley et al., 2015 Molecular Biology and Evolution 32:239-243; u = 2.9*10^-9 (95% CI = 1.3*10^-9 5.5*10^-9)


# Define functions for the inbreeding and maintenance design as well as 
# sequencing.
sequencing<-function(x,draws){
  # Pooled sequencing (to target coverage 40x)
  # x is now the frequency in the tube at the time of sequencing
  # read-depth is poisson, with a mean of 50x
  rpois(draws,50)->depth
  rbinom(draws,depth,x)/depth->allele_frequency_in_reads
  #return the frequency after the experiment and sequencing
  return(allele_frequency_in_reads)
}

inbreeding_design_and_sequencing<-function(x,draws){
  # x = the allele frequency among offspring of field caught females
  # they have mated once.
  
  # draws = the number of SNPs to draw
  
  # 8 generations of inbreeding
  # assuming free recombination! and 
  # randomly mating males and females (with equal allele frequencies)
  # (bad - I suspect this is anti-conservative, as LD will cause the observations to be more widely dispersed than this?
  # AW: LD will cause allele frequencies to be correlated (not sure about "more widely dispersed").
  for(i in 1:8){
    rbinom(draws,4,x)/4->x
  }
  
  # 40 generations of maintenance in the lab
  # Ne=40 (upper limit for an isofemale line?)
  # What if Ne = 100
  for(i in 1:40){
    rbinom(draws,80,x)/80->x
  }
  x<-sequencing(x,draws)
  return(x)
}


# Simulation parameters
# Simulate XX draws (nr. snps) of n individuals from a population.
# do this XXXX times [sims]
sims<-10
draws<-1000000 # nr. of snps
nes<-c(1000000,2000000,3000000,4000000) # Range of Ne
ns<-c(2,3,4,5,7,8,9,10) # Number of HIGH-LOW comparisons
MU <-c((1.0*10^-9),(2.8*10^-9),(2.9*10^-9),(5.5*10^-9),(8.4*10^-9)) # Range of mutation rates

# Comput the product 4*Ne*mu
alphas<-unique(4*expand.grid(nes,MU)[,1]*expand.grid(nes,MU)[,2])

#alphas<-0.022
#ns<-c(2,3,5,7,9)
field_fixed_diffs<-vector(length=sims*length(ns)*length(alphas))
fixed_diffs<-vector(length=sims*length(ns)*length(alphas))
Ns<-vector(length=sims*length(ns)*length(alphas))
Alphas<-vector(length=sims*length(ns)*length(alphas))
i<-1

for(alph in alphas){
    # Charlesworth and Charlesworth 2008 Elements of Population Genetics
    # This distribution describes a population at mutation-drift balance
    bet<-alph # alpha and beta are the same
    SFS<-rbeta(100000,alph,bet) # Define an allele frequency distribution
    # AW: a single SFS for "wild" population (rather than different for each population of origin)
    for(n in ns){
      sim<-1
      while(sim<=sims){
        cat("4*Ne*mu: ",alph," - N: ",n," - sim:",sim, " - i:",i,"\n")
        # Store the parameters
        Ns[i]<-paste("n = ",n,sep="")
        Alphas[i]<-paste("4*Ne*mu = ",alph,sep="")
        # Make matrices: 
        # ncols = nr populations
        # nrows = nr snps
        high_ps<-matrix(nrow = draws,ncol = n)
        low_ps<-matrix(nrow = draws,ncol = n)

        # Sample SNPs frequencies from the SFS 
        # in the offspring of two mated individuals.

        # HIGH lines
        high_ps<-apply(high_ps,2,
                       function(x){rbinom(n=draws,size=4,prob=SFS)/4})
        # LOW lines
        low_ps<-apply(low_ps,2,
                      function(x){rbinom(n=draws,size=4,prob=SFS)/4})
        # Sequencing
        high_ps<-apply(high_ps,2,
                       sequencing,draws=draws)
        low_ps<-apply(low_ps,2,
                      sequencing,draws=draws)
        
        # high_ps and low_ps are now the allele frequencies 
        # among the offspring of the field-sampled female.
        freqs<-cbind(high_ps,low_ps)

        # Remove all sites where the allele frequeny difference is 0 
        # (i.e. where all lines are fixed for the same allele)
        # Only interested in those that are variable, seen to vary.
        #freqs<-freqs[which(apply(freqs,1,function(x){!(all(x==1)|all(x==0))})),]
        high_ps<-freqs[,1:(ncol(freqs)/2)]
        low_ps<-freqs[,((ncol(freqs)/2)+1):ncol(freqs)]
        draw_diffs<-high_ps-low_ps
        
        # Find all sites where the allele frequency difference is 1
        diffs_tab<-table(apply(draw_diffs,1,
                               function(x){all(x==1)|all(x==-1)}))
        
        if(is.na(diffs_tab['TRUE'])){
          field_fixed_diffs[i]<-0
        }else{
          # Calculate the percent of consistently fixed differences in lines from the field
          field_fixed_diffs[i]<-(diffs_tab['TRUE']/sum(diffs_tab))*100
        }
        
        # Now run the inbreeding design and sequencing
        high_ps<-apply(high_ps,2,
                       inbreeding_design_and_sequencing,draws=draws)
        low_ps<-apply(low_ps,2,
                      inbreeding_design_and_sequencing,draws=draws)
        freqs<-cbind(high_ps,low_ps)
        
        # Only interested in those that are variable, i.e. that are seen to vary in the sample.
        freqs<-freqs[which(apply(freqs,1,function(x){!(all(x==1)|all(x==0))})),]
        high_ps<-freqs[,1:(ncol(freqs)/2)]
        low_ps<-freqs[,((ncol(freqs)/2)+1):ncol(freqs)]
        draw_diffs<-high_ps-low_ps
        diffs_tab<-table(apply(draw_diffs,1,
                               function(x){all(x==1)|all(x==-1)}))
        if(is.na(diffs_tab['TRUE'])){
          fixed_diffs[i]<-0
        }else{
          # Calculate the percent of consistently fixed differences in lines after inbreeding
          fixed_diffs[i]<-(diffs_tab['TRUE']/sum(diffs_tab))*100
        }
        sim<-sim+1
        i<-i+1
      }
    }
}

p_dat<-data.frame(field_diffs=field_fixed_diffs,inbr_diffs=fixed_diffs,
                  n=Ns,alpha=Alphas)
#p_dat
# Order the x-axis
# unique(p_dat$n)

p_dat$n<-factor(p_dat$n,
                levels = c("n = 2","n = 3","n = 4","n = 5",
                           "n = 7","n = 8","n = 9","n = 10"))
p_dat$alpha<-factor(as.character(p_dat$alpha))
# str(p_dat)

# Save data
write.table(p_dat,"~/Dropbox/st_andrews/pseudoobscura_wild_lines_MS/data/pseudo_wild_sims.csv",
            quote = FALSE,col.names = TRUE,row.names = FALSE,sep=",")

p_dat<-read.table("~/Dropbox/st_andrews/pseudoobscura_wild_lines_MS/BioRxiv_submission/scripts_data/pseudo_wild_sims.csv",
                  header=TRUE,sep=",")
p_dat$n<-factor(p_dat$n,
                levels = c("n = 2","n = 3","n = 4","n = 5",
                           "n = 7","n = 8","n = 9","n = 10"))
str(p_dat)
p_dat$alpha<-gsub("4\\*Ne\\*mu = ","",p_dat$alpha)
p_dat$alpha<-as.numeric(p_dat$alpha)
p_dat$alpha2<-as.factor(p_dat$alpha)
head(p_dat)
#str(p_dat)
# What percentage of the time do we expect all line comparisons
# to be fixed for the alternative?
tapply(p_dat$inbr_diffs,INDEX = list(p_dat$n),mean)
# What is the 95th percentile of the distribution of number of expected fixations
tapply(p_dat$inbr_diffs,INDEX = list(p_dat$n),quantile, probs=0.95)
tapply(p_dat$inbr_diffs,INDEX = list(p_dat$n),quantile, probs=0.05)
tapply(p_dat$inbr_diffs,INDEX = list(p_dat$n),quantile, probs=0.99)
# tapply(p_dat$inbr_diffs,INDEX = list(p_dat$n),max)
# 
p1<-ggplot()+
      geom_boxplot(data=p_dat,aes(n,inbr_diffs,colour=alpha,fill=alpha,group=interaction(n,alpha2)),
               position=position_dodge(0.9))+
  geom_hline(yintercept = 0.007,linetype="dashed",size=0.5)+
  xlab("")+
  scale_fill_gradient("4*Ne*mu",low="#fed976",high="#800026")+
  scale_colour_gradient("4*Ne*mu",low="#fed976",high="#800026")+
   ylab("Proportion of All\nFixed Differences (%)")+
   ggtitle("After Inbreeding\nand Sequencing is Simulated")+
   my.theme+
   theme(axis.text.x = element_text(size=14,face="bold",angle=45,hjust=1),
         axis.text.y = element_text(size = 14,face = "bold"),
         strip.text.y = element_text(size = 14,angle = 0),
         axis.title.y = element_text(size=14,face="bold"),
         legend.position = "right",
         legend.key = element_rect(fill="white"),
         legend.text = element_text(size=14,face="bold"),
         legend.title = element_text(size=14,face="bold"))
#
p2<-ggplot()+
   geom_boxplot(data=p_dat,aes(n,field_diffs,colour=alpha,fill=alpha,group=interaction(n,alpha2)),
                position=position_dodge(0.9))+
   geom_hline(yintercept = 0.007,linetype="dashed",size=0.5)+
   xlab("")+
  scale_fill_gradient("4*Ne*mu",low="#fed976",high="#800026")+
  scale_colour_gradient("4*Ne*mu",low="#fed976",high="#800026")+
  ylab("Proportion of All\nFixed Differences (%)")+
   ggtitle("Before Inbreeding\nand Sequencing is Simulated")+
   my.theme+
   theme(axis.text.x = element_text(size=14,face="bold",angle=45,hjust=1),
         axis.text.y = element_text(size = 14,face = "bold"),
         strip.text.y = element_text(size = 14,angle = 0),
         axis.title.y = element_text(size=14,face="bold"),
         legend.position = "right",
         legend.key = element_rect(fill="white"),
         legend.text = element_text(size=14,face="bold"),
         legend.title = element_text(size=14,face="bold"))
p2
library(cowplot)
cowplot::plot_grid(p2,p1,ncol=2)
```
