## Supplementary Text, Figures, and Table Legends for "Identifying genomic markers associated with female re-mating rate in *Drosophila pseudoobscura* by replicated bulk segregant analysis"

**Wiberg et. al.,**

### Supplementary Material

**Bioinformatic Pipeline**

In all steps outlined below, unless otherwise stated the default parameter values were used. See also the data associated with this study in the Dryad data repository (**LINK**)

**1) Trimming**

Trimming of the reads was carried out using Trimmomatic (v. 0.32; Bolger et al., 2014)

```
$ java -jar trimmomatic
	PE -phred33
	*R1_001.fastq.gz *R2_001.fastq.gz
	*tqc_R1_pe.fq.gz *ftqc_R1_se.fq.gz
	*tqc_R2_pe.fq.gz *ftqc_R2_se.fq.gz
	MINLEN:20 ILLUMINACLIP:${adapters}/TruSeq3-PE.fa:2:30:10
	SLIDINGWINDOW:1:20 MINLEN:20
```

**2) Mapping**

Mapping, indel re-alignment, removal of PCR duplicates and alignment filtering was carried out with bwa mem (v.; Li et al., 2009; Li, 2013) samtools (v. 1.2; Li et al., 2009), GATK (v. 3.3 McKenna et al., 2010; DePristo et al., 2011) and Picard (v. 2.14.1; Broad Institute).

**2.1) Mapping**

```
$ bwa mem -t 5 dpse-all-chromosome-r3.1.fasta *tqc_R1_pe.fq.gz
*tqc_R2_pe.fq.gz > *.sam
$ samtools view -Sb -q 20 *.sam > *.bam
$ samtools sort -@ 5 -o *_srt.bam *.bam
$ samtools rmdup *_srt.bam *_srt_rmdup.bam
$ samtools sort -@ 5 -o *_srt_rmdup_srt.bam *_srt_rmdup.bam
```

**2.2) Re-alignment around indels**

```
# Add random "readgroup" to .bam
$ java -jar $picard/AddOrReplaceReadGroups.jar
	I= *_srt_rmdup_srt.bam
	O= *_srt_rmdup_srt_rdgrp.bam
	RGID=1
	RGLB=L1
	RGPL=illumina
	RGPU=NONE
	RGSM=[READGROUPNAME]
# Index .bam file
$ samtools index *_srt_rmdup_srt_rdgrp.bam
# Re-align around indels
$ java -Xmx2g -jar $gatk
	-T RealignerTargetCreator
	-R dpse-all-chromosome-r3.1.fasta
	-I *_srt_rmdup_srt_rdgrp.bam
	-o *_srt_rmdup_srt_rdgrp.intervals
```

```
$ java -Xmx4g -jar $gatk
      -T IndelRealigner
      -R dpse-all-chromosome-r3.1.fasta
      -I *_srt_rmdup_srt_rdgrp.bam
      -targetIntervals *_srt_rmdup_srt_rdgrp.intervals
      -o *_srt_rmdup_srt_indraln.bam
# Sort reads and index
$ samtools sort -@ 5 -o *_srt_rmdup_srt_indraln_srt.bam
*_srt_rmdup_srt_indraln.bam
$ samtools index *_srt_rmdup_srt_indraln_srt.bam
```

**2.3) Coverage stats**

```
$ genomeCoverageBed -ibam *_srt_rmdup_srt_indraln_srt.bam >
*_srt_rmdup_srt_indraln_srt.cov
```

**3) SNP Calling**

SNP calling was performed with samtools mpileup (v. 1.2; Li et al., 2009), and files were converted to the .sync format with PoPoolation2 (v. 1.201; Kofler et al., 2011).

**3.1) Mpileup**

```
$ samtools mpileup
      -d 1500000
      -l
      -f dpse-all-chromosome-r3.1.fasta
      SLOB7.bam
      SLOC9.bam
      LEW17.bam
      LEW23.bam
      SHAA10.bam
      SHAC1.bam > *.mpileup
```

**3.2) Poopoolation2**

```
$ java -ea -Xmx20G
      -jar mpi2sync
      --input *.mpileup
      --output *.sync
      --fastq-type sanger
      --min-qual 20
      --threads 5
```

**4) Functional Analysis**

Functional analysis was carried out with GOwinda (Kofler & Schlötterer 2012) and the AME tool from the MEME package (Bailey et al., 2009; McLeay & Bailey 2010).

**4.1) Gowinda**

```
$ java -Xmx4G -jar ~/bin/Gowinda-1.12.jar
      --snp-file *all_snps.tab
      --candidate-snp-file *fixed_snps.tab
      --annotation-file Dpse_genes_dmelnames.gtf
      --gene-set-file dmel_funcassociate_go_associations_mod.txt
      --output-file *GOwinda.out
```

```
--simulations 1000000
--gene-definition updownstream1000000
```

**4.2) MEME**

```
$ ame --oc ame_out --pvalue-report-threshold 1 --control
*all_SNPs_regions.fasta *SNPs_regions.fasta motifDataBase
```

### Supplementary Figures

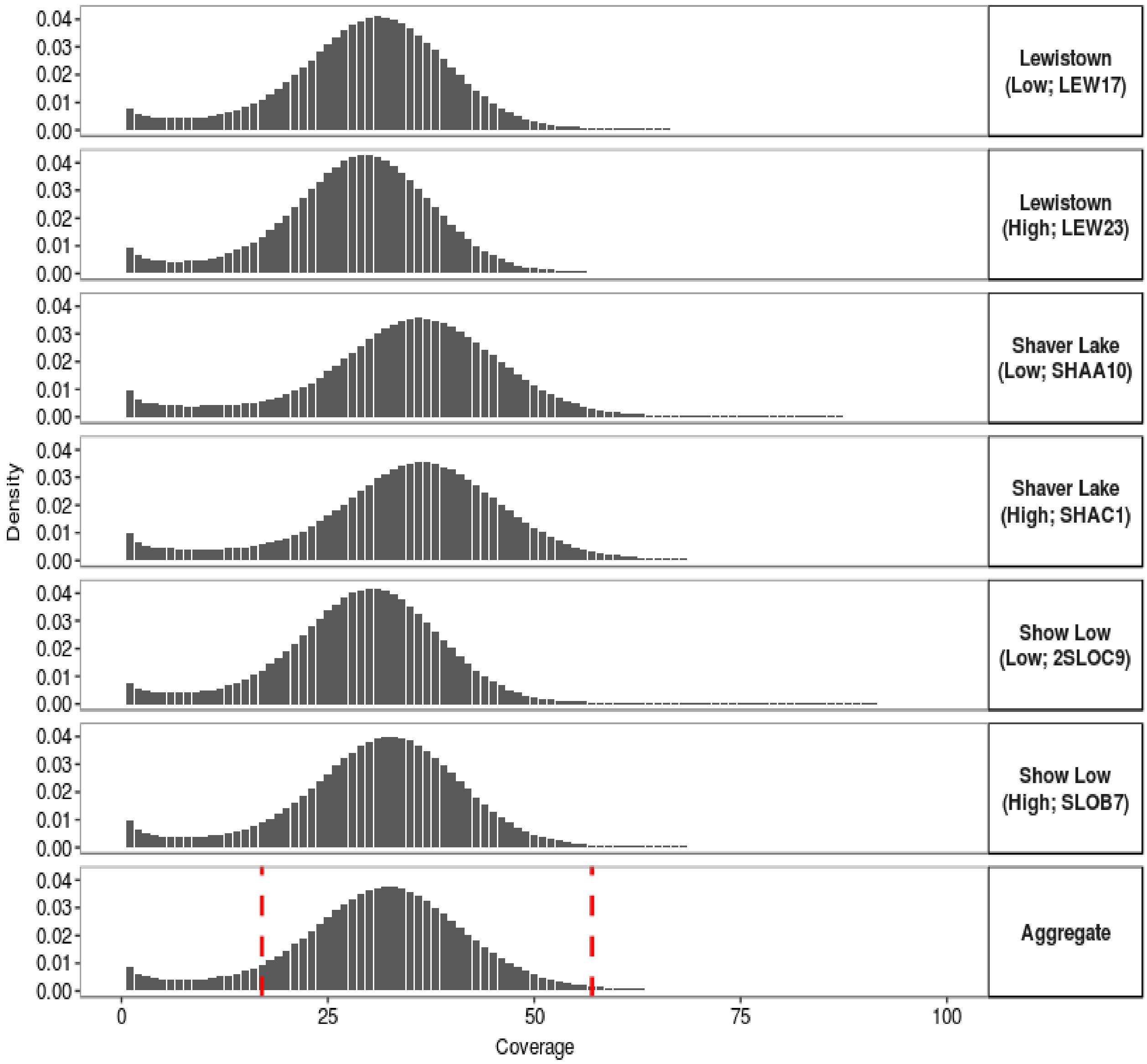


**Figure S1.** Coverage distributions across samples. Panels and panel labels refer to the isofemale lines in Table 1 and give the qualitative levels of female re-mating rates in each line. The "aggregate" distribution is obtained by summing counts in each coverage bin across samples. Red vertical lines give the $10^{th}$ and $90^{th}$ percentiles.

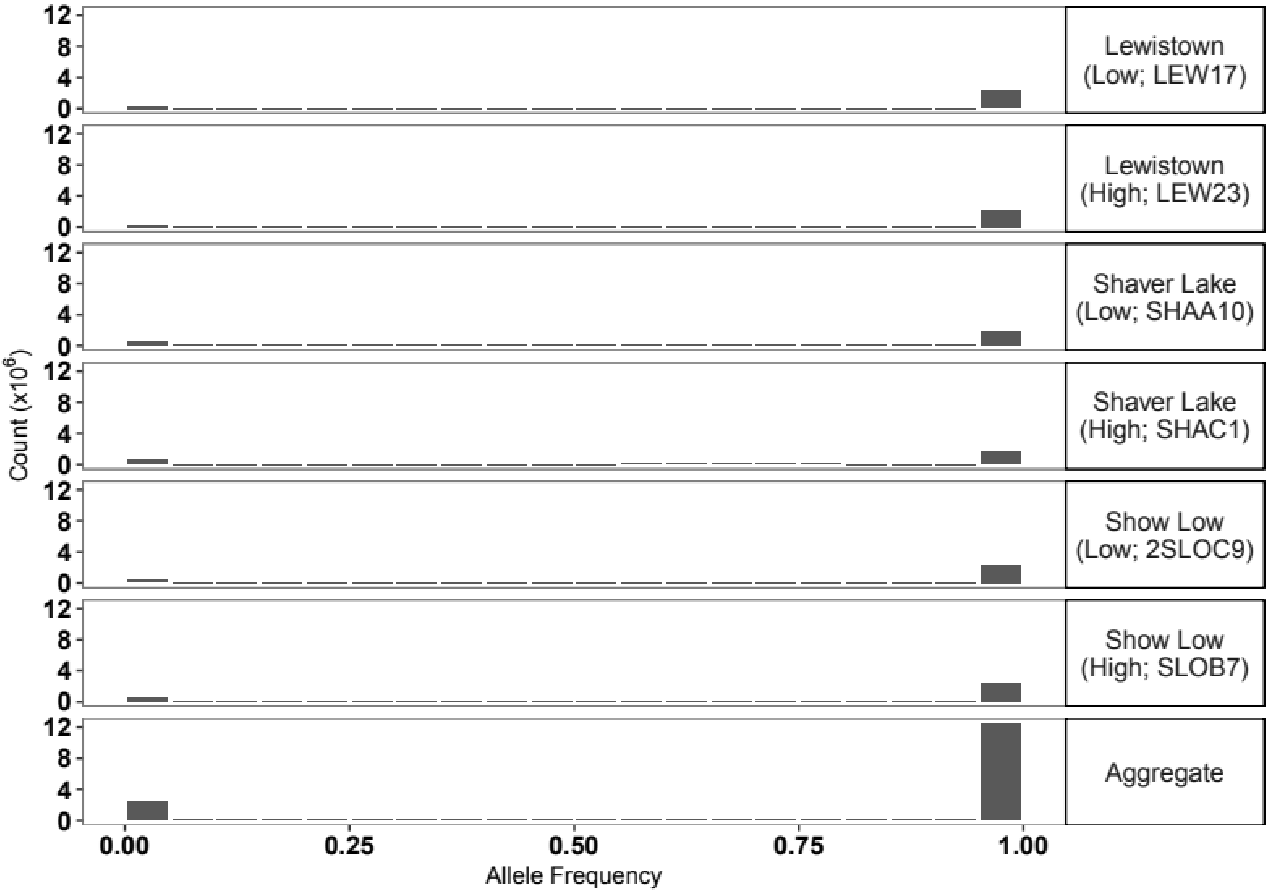


**Figure S2.** Allele frequency distributions of the overall major allele in all populations for **A)** all populations seprately, and **B)** summing counts from each bin across the populations into an aggregate distribution. Panel titles in **A)** give the population name, the level of female re-mating and the sample ID.

**Supplementary Tables and Legends**

**Table 1.** Summary statistics for populations and isofemale lines used in the current study. The population level re-mating rate is estimated from the proportion of clutches that show multiple paternity in Price et al., (2014).

| **Population** | **Isoline (Sample ID)** | **Isofemale line re-mating rate (%)** | **Population re-mating rate (%)** |
|---|---|---|---|
| Show Low | SLOB7 | 2.9 | 52 |
| | SLOC9 | 57 | |
| Lewistown | LEW17 | 5.6 | 92 |
| | LEW23 | 35 | |
| Shaver Lake | SHAA10 | 22 | 22 |
| | SHAC1 | 47 | |

**Table S2.** Summary statistics of sequencing, quality filtering and mapping steps for each sample. Figures before removal of duplicate reads, indel re-alignment, and sub-sampling of SLOB7, SHAC1, and SHAA10 samples are given in square brackets. The coverage distributions are shown graphically in figure 1.

| **Isoline (Sample ID)** | **Number of reads** | **% mapped (% proper pairs)** | **Mean coverage** |
|---|---|---|---|
| SLOC9 | ~51.02 m [~51 m] | 100 (97.92) [100 (97.94)] | 36.52x [37.08x] |
| SLOB7 | ~78.09 m [~79 m] | 100 (97.89) [100 (97.92)] | 34.49x [55.51x] |
| LEW17 | ~49.07 m [~49 m] | 100 (97.93) [100 (97.95)] | 35.16x [35.71x] |
| LEW23 | ~47.0 m [~47 m] | 100 (97.94) [100 (97.95)] | 33.83x [34.83x] |
| SHAC1 | ~47 m [~151.2 m] | 100 (95.3) [100 (95.30) ] | 42.38x [124.5x ] |
| SHAA10 | ~47 m [~109.6 m] | 100 (96.30) [100 (96.30)] | 42.11x [92.0x ] |

**Table S3.** Results for GO term enrichment analysis using GOwinda (v1.12; Kofler & Schlötterer, 2012). The table includes columns with the GO term ID, the average number of genes found per simulation, the number of genes associated with a SNP, the p-value, FDR corrected p-value, the number of unique genes associated with the SNP, the maximum possible number of genes which could be associated with a SNP, the number of genes in the GO term category, the GO term category description, a list of the genes associated with a SNP. Table includes results for genes within 1Mb of fixed SNPs.

**Table S4.** Fixed SNPs and the closest gene (within 1Mb) from the *D. pseudoobscura* annotation. The table includes the flybase gene ID, the annotation symbol, the gene name, and the gene annotation symbol.

**Table S5.** Tables of results from modPhEA. The table includes columns for each tested phenotypic class, a descriptive name for each class, the proportion of genes near fixed SNPs which are annotated with that phenotypic class, the proportion of genes from the rest of the genome which are annotated with that phenotypic class, p-value from a Fisher's exact test of association, a Benjamini-Hochberg adjusted p-value, and a Bonferroni adjusted p-value. The table includes results for genes within 1Mb of fixed SNPs.

**Table S6.** Genes near fixed SNPs which are also in the phenotypic class "behaviour defective." The table includes the flybase gene ID, the annotation symbol, the gene name, and the gene annotation symbol. The table includes results for genes within 1Mb of fixed SNPs.

**Table S7.** Results for AME transcription factor (TF) motif enrichment analysis. Threshold p-value for reporting results was set to 1000 (i.e. all results are reported for completeness). The table includes the names of all motifs tested for enrichment, the p-value from a Fisher's exact test, and the p-value after correction for multiple testing (by Bonferroni correction).
